# Which evolutionary game-theoretic model best captures NSCLC dynamics?

**DOI:** 10.1101/2025.07.10.664060

**Authors:** Hasti Garjani, Johan Dubbeldam, Kateřina Staňková, Joel S. Brown

**Affiliations:** Delft Institute of Applied Mathematics, Delft University of Technology, Delft, The Netherlands; Institute for Health Systems Science, Delft University of Technology, Delft, The Netherlands; Integrated Mathematical Oncology, H. Lee Moffitt Cancer and Research Institute, Tampa, FL, United States of America

## Abstract

Understanding and predicting the eco-evolutionary dynamics of cancer requires identifying mathematical models that best capture tumor growth and treatment response. In this study, we fit a family of two-population models to *in-vitro* data from non-small cell lung cancer (NSCLC), tracking drug-sensitive and drug-resistant cells under varying environmental conditions. The dataset, originally presented by Kaznatcheev et al., includes conditions with and without the drug Alectinib and cancer-associated fibroblasts (CAFs). We compare combinations of growth models (logistic, Gompertz, von Bertalanffy) and drug efficacy terms (Norton–Simon, linear, ratio-dependent) to identify which best explains the observed dynamics. Our models incorporate density dependence, frequency-dependent competition, and drug response, enabling mechanistic interpretation of tumor cell interactions. The logistic model with ratio-dependent drug efficacy best fits monoculture data. Using growth parameters from monocultures, we infer inter-type competition coefficients in co-cultures. We find that growth rate and carrying capacity are stable across CAF conditions, while competition and drug efficacy parameters shift, altering interaction dynamics. Notably, CAFs promote coexistence between resistant and sensitive cells, whereas Alectinib induces competitive exclusion. Our results underscore the need to evaluate both model fit and biological plausibility to guide therapeutic modeling of cancer.

**Author summary:** How cancer cells grow, compete, and respond to treatment depends not only on the drug, but also on their ecological context, including interactions with other cells and components of the tumor microenvironment. In this study, we explore how different mathematical models capture the behavior of non-small cell lung cancer (NSCLC) cells under various conditions. We focus on two cell populations: one sensitive to treatment, and one resistant. Using *in-vitro* data, we compare growth models and drug response types to identify which model best explains the observed population dynamics. We also investigate how different factors, such as the drug Alectinib and cancer-associated fibroblasts (CAFs), change the way cancer cells interact. Our game-theoretic approach allows us to quantify how these external conditions affect competition between cell types, revealing when resistant and sensitive cells can coexist. These findings contribute to a deeper understanding of tumor ecology and may support the development of adaptive cancer therapies that anticipate evolutionary responses to treatment.

## Introduction

Lung cancer is one of the most common cancers diagnosed [1], and it is the leading cause of cancer death [2]. Despite advances in treating cancer, cures for metastatic cancers are rare [3–5]. Mathematical oncology provides mathematical models that can help us understand the response of cancer cells to therapy and improve anti-cancer therapies [6, 7].

Increasingly, cancer progression and growth are viewed as an evolutionary game where the best strategy (heritable trait) of an individual cancer cell depends upon the strategies of its neighboring cancer cells and the therapy [8–11]. We can also assume cancer is involved in a leader-follower (Stackelberg) evolutionary game where the physician (leader) applies therapy to which the cancer cells (followers) respond [12–14]. Cancer cells’ interactions with their environment include competition for resources, co-opting of normal cells, responding to immune cells, and engaging in public goods games through angiogenesis and ecosystem engineering [15–17].

Competition within evolutionary games can be driven by three types of processes [18, 19]. First, there can be density-independent processes where a cancer cell’s survival and proliferation rate are (nearly) independent of other cells – this is typical of population growth at low population sizes when space and nutrients are abundant. Second, density-dependent processes. These occur when a cancer cell suffers lower fitness (prospects for survival and proliferation) as the density of cancer cells increases – typically occurring through lack of space or nutrients or build-up of toxins. Third, there are frequency-dependent interactions which occur when the fitness of a cell depends on its trait and the frequency of traits among the other cancer cells [20].

In oncologic literature, there is growing interest in competition assays and experiments where two (or sometimes more) cell lines are followed in mono- and co-cultures to see how the populations grow, interact, and do or do not coexist. Such experiments draw from a long tradition in ecology of competing single-celled protists such as Paramecium or yeast. This tradition goes all the way back to the seminal experiments of Gause in the 1920s and 30s [21]. For cancer, such experiments can reveal the cost of resistance (competing a sensitive and resistant cell line under the presence or absence of drug therapy) [22], the role of glycolysis (competing highly glycolytic cell lines against highly oxidative phosphorylation cell lines) [23], effects of immune cells or fibroblasts (competing cell lines thought to differ in the costs and benefits associated with normal cells), effects of microenvironmental conditions (competing cells lines with differing responses to pH, nutrient deprivation, growth factors, etc.) [24–26], effects of different therapies singly or in combination (competing cell lines through having differing responses), and effects of gene knock-out experiments to discern metabolic pathways (competing knock-out cell lines against their sensitive cell line) [27].

Considering the interactions between the cells as a game, there are roughly three ways to model the trajectory and outcome of competition experiments involving two cell lines. The simplest is to focus entirely on frequency-dependent interactions by using a 2 × 2 matrix game where the cell types represent two strategies, and they experience payoffs associated with intra-cell line and inter-cell line interactions. The outcomes can be determined by estimating entries of the payoff matrix and using the replicator equation to derive dynamics towards an equilibrium [28]. This approach subsumes any density-independent and density-dependent processes into the 4 elements of the payoff matrix. A more sophisticated version of the 2 × 2 matrix game occurs when the game is no longer bilinear, meaning that the success of an individual is no longer a linear weighted averaging of payoffs based on the two cell lines. Rather, payoffs can be a nonlinear (concave up or concave down) combination of payoffs from changing the frequency of cell types [29]. To model the trajectory and outcomes, one requires fitness functions based on these nonlinear relationships that can be used to directly derive frequency dynamics. Such a frequency-dependent model will produce trajectories that deviate somewhat from the replicator equation based on the degree of nonlinearities. A third approach uses ecological models of competition, such as Lotka-Volterra (L-V) competition equations. Such a model can include density-independence, *ρ*, density dependence via a carrying capacity, *K*, and frequency-dependent interactions via the inter-type competition coefficients, *α*_*ij*_. The outcomes of and trajectories of such a model, like the L-V competition equations, are determined by these parameters. We use this model. Properly fitting such a model requires relatively large numbers of treatment combinations using initial conditions. Namely, co-cultures seeded at different frequencies of cell types and ideally at different total initial number of cells.

Here, we test for frequency- and density-dependent interactions between two cancer cell lines using data from [25]. They measured the population of two cell lines using 12 initial ratios from zero to one, under four environmental conditions: all combinations of the presence and absence of cancer-associated fibroblasts (CAFs) and the presence and absence of the drug. In [25], they fit their data to the replicator equation, which only considers the frequency dynamics of the two cell types, not the dynamics of their population sizes.

Here, we expand their analyses by explicitly modeling the population dynamics. First, we fit the data to three models of population growth (Logistic, Gompertz, Von Bertalanffy [30, 31]) that are commonly applied to cancer. We extend the Von Bertalanffy growth dynamics to consider competition between the two cell types in a manner similar to Logistic and Gompertz [32, 33]. Second, we compare three ways of including drug efficacy [34]: Norton-Simon model [35], density-independent mortality [36], and density-dependent mortality [37]. Third, we test how intrinsic growth rates (*ρ*s as density-independent effects), carrying capacities (*K*s as density-dependent effects), and competition coefficients (*α*_*ij*_s as frequency-dependent effects) vary with cell type, presence and absence of CAFs, and presence and absence of drug. Finally, we compare our model predictions for the outcomes of competition under the four environments to those of [25].

## Methods

Kaznatcheev et al. [25] conducted competition experiments in cell culture using two non-small cell lung cancer cell lines: one sensitive (parental) and the other resistant to the drug Alectinib. Experimental treatments consisted of all combinations of the presence or absence of cancer-associated fibroblasts (CAFs) and the presence or absence of Alectinib, and 8 seeding frequencies (0, 0.1, 0.2, 0.4, 0.6, 0.8, 0.9, and 1 ratios of sensitive to resistant cells). Each combination was replicated 6 times. To measure population dynamics, sensitive and resistant cell lines were labeled with green fluorescent protein and mCherry fluorescent proteins, respectively. Population sizes as measured by fluorescence intensity were recorded at intervals of 4 hours from time 0 to 120 hours. When present, Alectinib was introduced into wells as a single dose after 20 hours. Kaznatcheev et al. [25], in evaluating the outcomes of competition, only considered the frequency-dynamics of cells within the wells, namely the ratio of sensitive cells to total population size. In what follows, we use their entire time series of the cancer cells’ population sizes to fit models from population ecology. We investigate models that consider density independence, density dependence, and frequency dependence. We first evaluate how well the data fit three commonly used growth models of population dynamics, adjusted for two-species competitive interactions. Next, we determine the representative drug efficacy model. Then, focusing on the Lotka-Volterra competition models, we test how cell lines, fibroblasts, and the drug influence intrinsic growth rates, carrying capacities, and competition coefficients. Finally, we assess how these parameter variations impact interactions among different cell types.

### Logistic, Gompertz, and Von Bertalanffy growth models

We look into two-population extensions of the Logistic, Gompertz, and Von Bertalanffy models. The two population models with Logistic and Gompertz growth have been explored for cancer population growth in the literature [20, 33]. The general forms of these two-population models with Norton-Simon drug effects are presented in Eq 1 and 2, respectively.

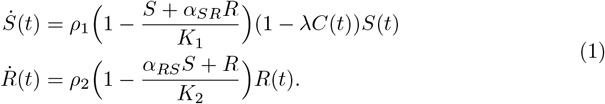

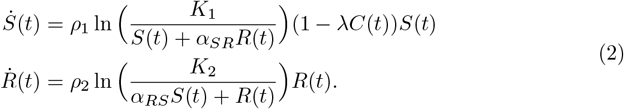

The three variables *S*(*t*), *R*(*t*), and *C*(*t*) show the sensitive to drug population, the resistant to drug population, and the amount of drug. We extend Von Bertalanffy growth dynamics [30] to a two-population model in Eq 3.

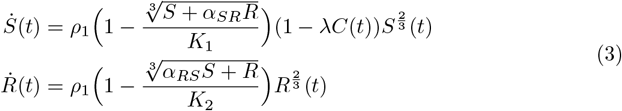

These models consider density independence through *ρ*_1_ and *ρ*_2_, density dependence through *K*_1_ and *K*_2_, frequency dependence through *α*_*SR*_ and *α*_*RS*_, and drug efficacy through *λ*. We fit the data to investigate how many and which of these parameters are supported by the dataset derived from the *in-vitro* experiment.

We fit each model to each repetition of the experiment individually to examine the variability in parameter estimates. The model simultaneously predicts values for resistant and sensitive cells, and given that, in most instances, one population type significantly outnumbers the other, we use a weighted sum of squared errors. The corresponding cost function for each well is denoted as:

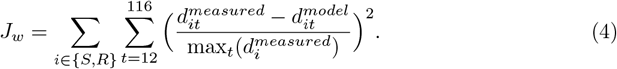

where *i* ∈ {*S, R*} denotes the population type, and *t* ∈ {12, 16, …, 116} is the time point in the series. The equation 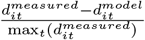, presents the error value for cell of type *i* at time *t*. We start from *t* = 12 hour to let the cells settle in the well as suggested by [25]. The 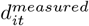 is the measured population at time *t* for population type *i*, 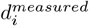 is the vector of the measured population of type *i* from time 12 to 116 (measured every four hours), and 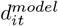 is the predicted population size using the proposed model.

We use the Chi-square error (*χ*^2^ = *J*_*w*_) and AIC (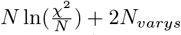 where *N*_*varys*_ is the number of variables, and *N* is the number of data points) as our goodness of fit measures to compare the fit results in different wells.

We estimate the model parameters by minimizing the cost function Eq 4, considering the difference between the predicted data points from time 12 hours until time 116 hours and the measured data points. To fit the data to the two-population models, the initial values for the resistant and sensitive populations are set to the measured data values at 12 hours, as Kaznatcheev et al. recommended in their paper. Here, we fit the observations in each well to the model separately to ensure that the intra-subject variations are not averaged out. This helps in assessing whether significant variations occur because of differing experimental conditions, such as varying initial ratios or initial populations

These models include a maximum of seven different parameters (*ρ*_1_, *ρ*_2_, *K*_1_, *K*_2_, *α*_*SR*_, *α*_*RS*_, and *λ*). First, we fit the data to a model with two unknown parameters (same growth rates and carrying capacities for both cancer cell types) in cases where the drug is not present and a model with three unknown parameters (same growth rates and carrying capacities for both cancer cell types and drug efficacy for the sensitive population) when the drug is present. We increase the number of unknown parameters to at most 5, as seen in Table 1. To prevent overfitting, we increase the number of parameters gradually, considering equal *ρ*_1_ and *ρ*_2_ values, equal *K*_1_ and *K*_2_ values, and *α*_*SR*_ = *α*_*RS*_ = 1. We assume distinct growth rates for the two populations *ρ*_1_ ≠ *ρ*_2_ for the second fit (Logistic 2 of Table 1). For the third and fourth fits, we assume that the model has the same growth rates and one competition coefficient equal to one and an unknown *α*_*SR*_ or *α*_*RS*_, respectively. Finally, we fit the model with distinct competition coefficients and the same growth rates and carrying capacities for both populations.

**Table 1.** Models inspired by Logistic growth.

|  |  |
| --- | --- |
| Logistic 1 | $\dot{S}(t) = \rho \left( 1 - \frac{S+R}{K} \right) (1 - \lambda C(t)) S(t)$ $\dot{R}(t) = \rho \left( 1 - \frac{S+R}{K} \right) R(t).$ |
| Logistic 2 | $\dot{S}(t) = \rho_1 \left( 1 - \frac{S+R}{K} \right) (1 - \lambda C(t)) S(t)$ $\dot{R}(t) = \rho_2 \left( 1 - \frac{S+R}{K} \right) R(t).$ |
| Logistic 3 | $\dot{S}(t) = \rho \left( 1 - \frac{S+\alpha_{SR}R}{K} \right) (1 - \lambda C(t)) S(t)$ $\dot{R}(t) = \rho \left( 1 - \frac{S+R}{K} \right) R(t).$ |
| Logistic 4 | $\dot{S}(t) = \rho \left( 1 - \frac{S+R}{K} \right) (1 - \lambda C(t)) S(t)$ $\dot{R}(t) = \rho \left( 1 - \frac{\alpha_{RS}S+R}{K} \right) R(t).$ |
| Logistic 5 | $\dot{S}(t) = \rho \left( 1 - \frac{S+\alpha_{SR}R}{K} \right) (1 - \lambda C(t)) S(t)$ $\dot{R}(t) = \rho \left( 1 - \frac{\alpha_{RS}S+R}{K} \right) R(t).$ |

Sensitive and resistant population growth follows the ODE model presented at each block. The number of unknown parameters in the Logistic 1 model are two or three depending on whether the drug is present or not ( $\rho, K$ , and $\lambda$ ). Unknown parameters in Logistic 2 are $\rho_1, \rho_2, K$ , and $\lambda$ . Unknown parameters in Logistic 3 are $\rho, K, \alpha_{SR}$ , and $\lambda$ . Unknown parameters in Logistic 4 are $\rho, K, \alpha_{RS}$ , and $\lambda$ . Unknown parameters in Logistic 5 are $\rho, K, \alpha_{SR}, \alpha_{RS}$ , and $\lambda$ .

The models are illustrated in Table 1.

Sensitive and resistant population growth follows the ODE model presented at each block. The number of unknown parameters in the Logistic 1 model are two or three depending on whether the drug is present or not (*ρ, K*, and *λ*). Unknown parameters in Logistic 2 are *ρ*_1_, *ρ*_2_, *K*, and *λ*. Unknown parameters in Logistic 3 are *ρ, K, α_SR_*, and *λ*. Unknown parameters in Logistic 4 are *ρ, K, α_RS_*, and *λ*. Unknown parameters in Logistic 5 are *ρ, K, α_SR_, α_RS_*, and *λ*.

We repeat the same stepwise increase in the number of unknown parameters for Eq 2 and Eq 3. The tables containing these models are in the appendix (Table 2 and Table 3). In total, we fit 36 wells to 15 different models. We compare the error and AIC distribution of the different models to determine the appropriate models that fit our data.

**Table 2.** Models based on Gompertz growth.

|  |  |
| --- | --- |
| Gompertz 1 | $\dot{S}(t) = \rho \ln\left(\frac{K}{S(t)+R(t)}\right)(1 - \lambda C(t))S(t)$<br>$\dot{R}(t) = \rho \ln\left(\frac{K}{S(t)+R(t)}\right)R(t).$ |
| Gompertz 2 | $\dot{S}(t) = \rho_1 \ln\left(\frac{K}{S(t)+R(t)}\right)(1 - \lambda C(t))S(t)$<br>$\dot{R}(t) = \rho_2 \ln\left(\frac{K}{S(t)+R(t)}\right)R(t).$ |
| Gompertz 3 | $\dot{S}(t) = \rho \ln\left(\frac{K}{S(t)+\alpha_{SR}R(t)}\right)(1 - \lambda C(t))S(t)$<br>$\dot{R}(t) = \rho \ln\left(\frac{K}{S(t)+R(t)}\right)R(t).$ |
| Gompertz 4 | $\dot{S}(t) = \rho \ln\left(\frac{K}{S(t)+R(t)}\right)(1 - \lambda C(t))S(t)$<br>$\dot{R}(t) = \rho \ln\left(\frac{K}{\alpha_{RS}S(t)+R(t)}\right)R(t).$ |
| Gompertz 5 | $\dot{S}(t) = \rho \ln\left(\frac{K}{S(t)+\alpha_{SR}R(t)}\right)(1 - \lambda C(t))S(t)$<br>$\dot{R}(t) = \rho \ln\left(\frac{K}{\alpha_{RS}S(t)+R(t)}\right)R(t).$ |

Sensitive and resistant population growth follows the ODE model presented at each block. The number of unknown parameters in the Gompertz 1 model is two or three, depending on whether the drug is present or not ( $\rho$ , $K$ , and $\lambda$ ). Unknown parameters in Gompertz 2 are $\rho_1$ , $\rho_2$ , $K$ , and $\lambda$ . Unknown parameters in Gompertz 3 are $\rho$ , $K$ , $\alpha_{SR}$ , and $\lambda$ . Unknown parameters in Gompertz 4 are $\rho$ , $K$ , $\alpha_{RS}$ , and $\lambda$ . Unknown parameters in Gompertz 5 are $\rho$ , $K$ , $\alpha_{SR}$ , $\alpha_{RS}$ , and $\lambda$ .

**Table 3.** Models based on Von Bertalanffy growth.

|  |  |
| --- | --- |
| Von Bertalanffy 1 | $\dot{S}(t) = \rho(1 - \frac{\sqrt[3]{S+R}}{K})(1 - \lambda C(t))S^{\frac{2}{3}}(t)$<br>$\dot{R}(t) = \rho(1 - \frac{\sqrt[3]{S+R}}{K})R^{\frac{2}{3}}(t).$ |
| Von Bertalanffy 2 | $\dot{S}(t) = \rho_1(1 - \frac{\sqrt[3]{S+R}}{K})(1 - \lambda C(t))S^{\frac{2}{3}}(t)$<br>$\dot{R}(t) = \rho_2(1 - \frac{\sqrt[3]{S+R}}{K})R^{\frac{2}{3}}(t).$ |
| Von Bertalanffy 3 | $\dot{S}(t) = \rho(1 - \frac{\sqrt[3]{S+\alpha_{SR}R}}{K})(1 - \lambda C(t))S^{\frac{2}{3}}(t)$<br>$\dot{R}(t) = \rho(1 - \frac{\sqrt[3]{S+R}}{K})R^{\frac{2}{3}}(t).$ |
| Von Bertalanffy 4 | $\dot{S}(t) = \rho(1 - \frac{\sqrt[3]{S+R}}{K})(1 - \lambda C(t))S^{\frac{2}{3}}(t)$<br>$\dot{R}(t) = \rho(1 - \frac{\sqrt[3]{\alpha_{RS}S+R}}{K})R^{\frac{2}{3}}(t).$ |
| Von Bertalanffy 5 | $\dot{S}(t) = \rho(1 - \frac{\sqrt[3]{S+\alpha_{SR}R}}{K})(1 - \lambda C(t))S^{\frac{2}{3}}(t)$<br>$\dot{R}(t) = \rho(1 - \frac{\sqrt[3]{\alpha_{RS}S+R}}{K})R^{\frac{2}{3}}(t).$ |

Sensitive and resistant population growth follows the ODE model presented at each block. The number of unknown parameters in the Von Bertalanffy 1 model is two or three, depending on whether the drug is present or not ( $\rho$ , $K$ , and $\lambda$ ). Unknown parameters in Von Bertalanffy 2 are $\rho_1$ , $\rho_2$ , $K$ , and $\lambda$ . Unknown parameters in Von Bertalanffy 3 are $\rho$ , $K$ , $\alpha_{SR}$ , and $\lambda$ . Unknown parameters in Von Bertalanffy 4 are $\rho$ , $K$ , $\alpha_{RS}$ , and $\lambda$ . Unknown parameters in Von Bertalanffy 5 are $\rho$ , $K$ , $\alpha_{SR}$ , $\alpha_{RS}$ , and $\lambda$ .

### Drug efficacy in the two-population model

In addition to growth dynamics, we analyze three models of drug efficacy: the Norton-Simon model, density-independent mortality, and density-dependent mortality. The three mentioned drug effects for the logistic model are illustrated in Eqs 5-7.

1. Norton-Simon model:

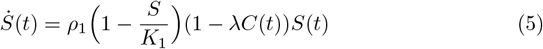
2. Linear drug effect:

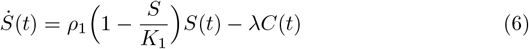
3. Ratio-dependent drug effect:

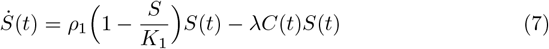

We determine the best drug efficacy model by fitting it to the monoculture data. Evidence suggests the model’s structural characteristics (e.g., carrying capacity values, the representative growth model, or drug efficacy) for monoculture and co-culture are similar [38]. Therefore, we use the parameter estimates of the monoculture to develop the two-population model.

### Impact of cell type and the presence or absence of CAFs and drug on model parameters

By fitting data from each well to the model separately, we get replicate parameter estimates, which allow us to compare within-category variance and between-category variance. We use ANOVA (Analysis of Variance) to test the impact of CAFs and therapy on growth rates (density-independent parameters *ρ*) and carrying capacities (density-dependent parameters *K*).

First, we find the growth rate and carrying capacity of monoculture-sensitive and resistant cells in DMSO and CAF environments. Next, we use a two-way ANOVA test for the effects of population type (sensitive and resistant), environment (DMSO and CAF), and their interaction. Then, we use these estimates to estimate the drug efficacy term when the drug is present. The fitted one-population model of sensitive cells:

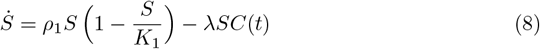

The fitted one-population model of resistant cells:

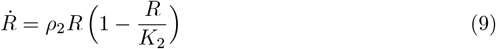

The co-culture experiment is also carried out in four distinct environments. We use the growth rate, carrying capacity, and drug effect parameters derived from the monoculture experiment to find the competition coefficients in the co-culture. This stepwise approach leverages our prior knowledge of the model in monocultures. The competition coefficients are the only variables that change while fitting the co-culture data.

### Effect of the environment on the interactions among the cells

The experiment is conducted under four environmental conditions, accounting for the presence or absence of CAFs and the presence or absence of the drug. Kaznatcheev et al. [25] only investigated frequency, not density dependent dynamics. They used the replicator equation to forecast outcomes. They predicted the occurrence of a mixed equilibrium (where both sensitive and resistant cells coexist) only in an environment with the presence of CAF, while anticipating the extinction of sensitive cells in all other environments. We fit the data to a model that incorporates density dependence, density independence, and frequency dependence. We examine the influence of the environment on each parameter based on the values of the fitted parameters to gain more insight into the effect of the environment.

## Results

### Comparing Logistic, Gompertz, and Von Bertalanffy growth models

Our analysis demonstrates that fitting the two-population Logistic, Gompertz, and Von Bertalanffy models outlined in Section results in satisfactory fits for all models with at least three fitted parameters. We find AIC and error distributions for specific environments or specific number of unknown parameters that suggest one growth model is better. Still, the superiority of each growth model has no consistency. This can be due to the high correlation among the parameters and similarities of the models. The median error of the 15 models fit is illustrated in Fig 1. As seen in this figure, the median error is large when we have only two unknown parameters, such as in the model referred to as “Logistic 1” in Table 1. However, the difference between the error values for the rest of the models is very small, as presented in Fig 1. Comparisons based on the AIC align with the conclusions drawn from the error metric, as illustrated in Appendix Fig 8.

**Fig 1.**
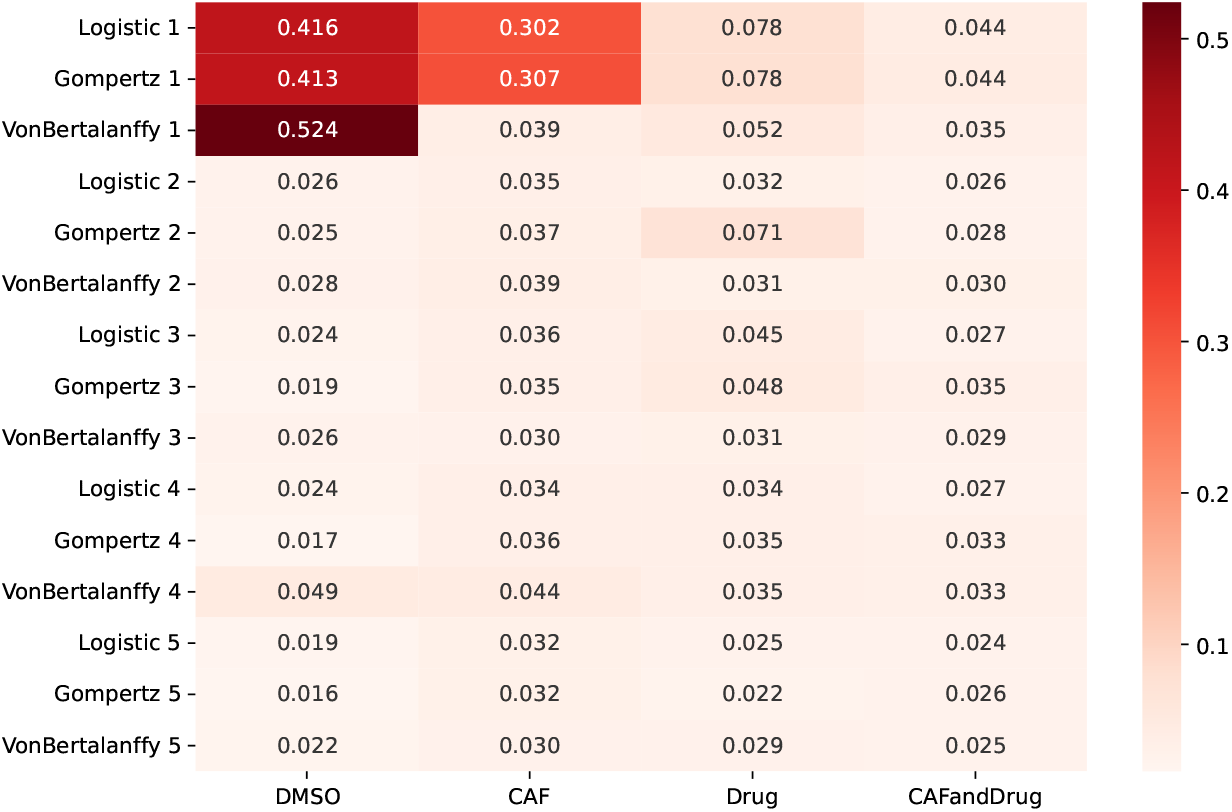
Chi-square error for the two-population model fits. Heatmap of the median of error values is for the proposed fifteen models in Tables 1, 2, 3 that account for Logistic, Gompertz, and Von Bertalanffy growth models with different number of unknown parameters. Different growth models with more than two parameters result in insignificant error variations.

Note that, allowing all the parameters to vary independently for each well results in overfitting, shown by minimal variations in error values as the number of parameters increases. There is insufficient variation in goodness of fit metrics for determining the best growth model. Therefore, we fitted the logistic model as it makes no assumptions regarding nonlinear density-dependence. Results from an ANCOVA provide additional support for this decision, as presented in the Appendix.

### Determining drug efficacy term

We analyzed three treatment effects: Norton-Simon, linear, and ratio-dependent drug efficacies, introduced in Eq 5, Eq 6, and Eq 7, respectively. Among these three models, ratio-dependent drug efficacy was more representative of the data and led to smaller chi-square errors, as observed in Fig 2. Furthermore, ratio-dependent drug efficacy aligns with the accepted biological understanding of population growth in relation to resource scarcity, whereas the Norton-Simon model does not. Our model uses *C*(*t*) = 1 when the drug is administered, since the drug is administered in a constant dose [25]. Therefore, in the Norton-Simon model, for any *λ >* 1, if the population exceeds the carrying capacity *K*, the population derivative 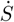 becomes positive and can increase indefinitely. Furthermore, for *λ <* 1, the model is not representative of a decreasing population, indicating that this model does not accurately reflect the results of this experiment, where certain wells exhibit population decline.

**Fig 2.**
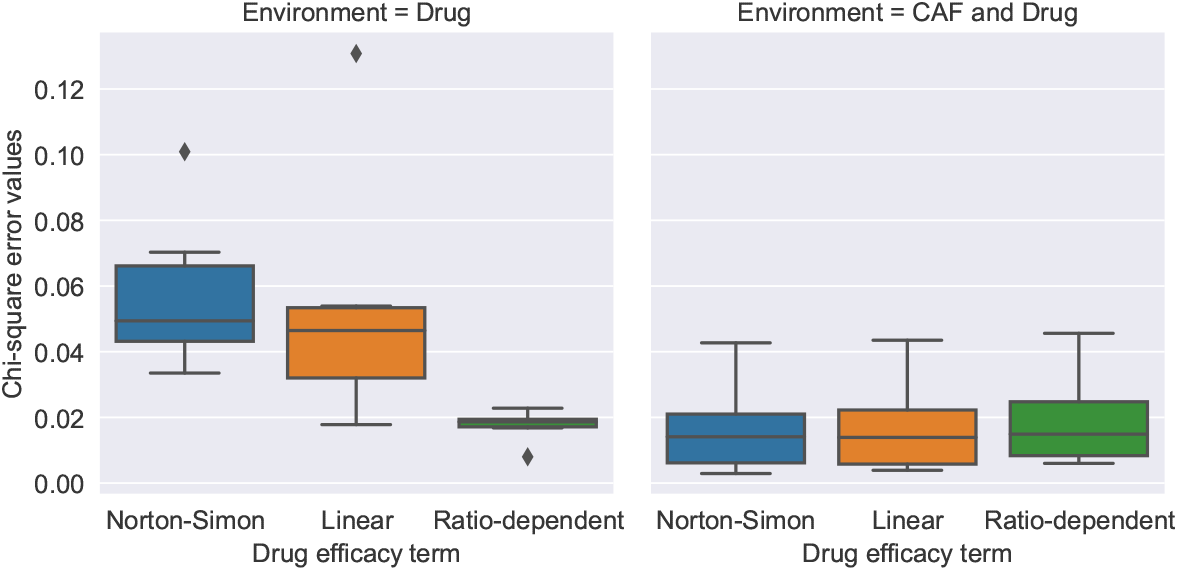
Chi-square error values for Norton-Simon, linear, and ratio-dependent drug efficacy (Eq 5, 6, 7) in the presence of drug. The ratio-dependent drug efficacy model leads to smaller errors compared to Norton-Simon and linear drug effects in environments when only the drug is present.

### Parameter variations across cell types and CAFs presence

We used the logistic growth and the ratio-dependent treatment effect determined in sections and to build the co-culture model. First, we fitted the sensitive and resistant cell population in wells where only one cell type is present (monocultures) in DMSO and CAF environments. We found estimates for growth rates of sensitive and resistant cells (*ρ*_1_ and *ρ*_2_) and their carrying capacities (*K*_1_ and *K*_2_). We observed that the growth rate and carrying capacity values of sensitive and resistant cells were not influenced by the presence and absence of CAFs when the drug was not present (Fig 3, Fig 4).

**Fig 3.**
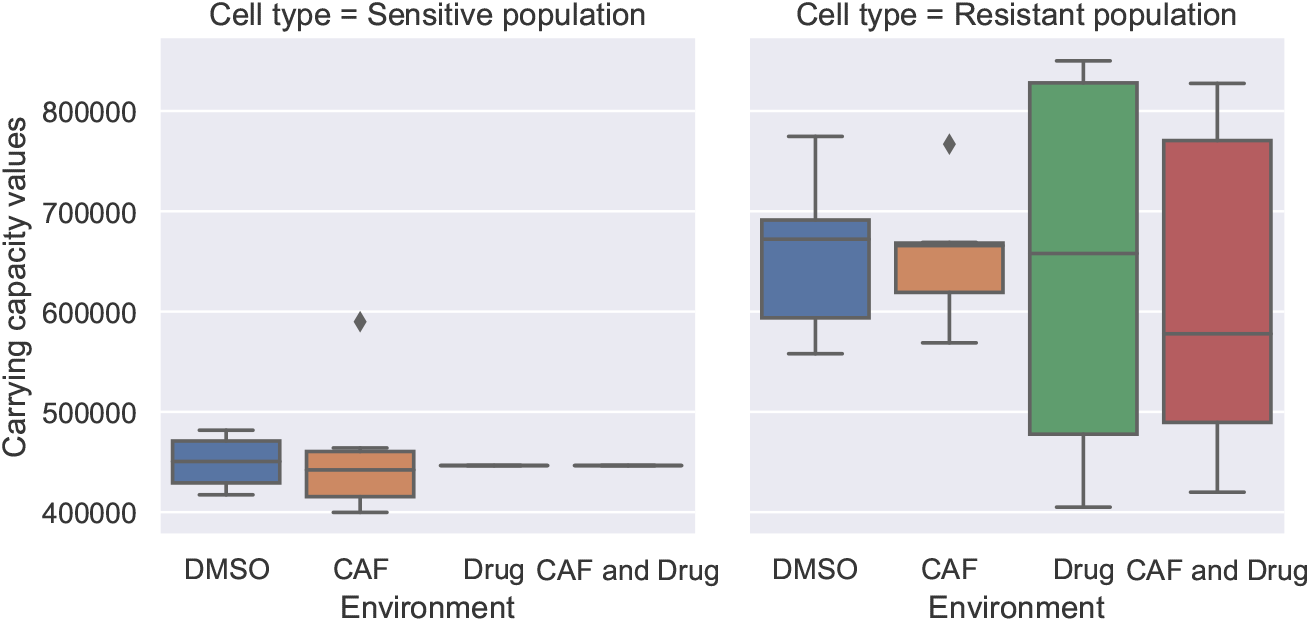
Carrying capacities (*K*_1_ and *K*_2_) in all environments. The boxplots of carrying capacity for sensitive and resistant populations illustrate the outcomes in different wells of the monotypic cultures. The carrying capacity of sensitive cells in the presence of the drug is fixed at the median of those parameters in DMSO and CAF environments. The cell type significantly affects carrying capacity, while the presence or absence of CAFs and drug results in minimal and non-significant variation.

**Fig 4.**
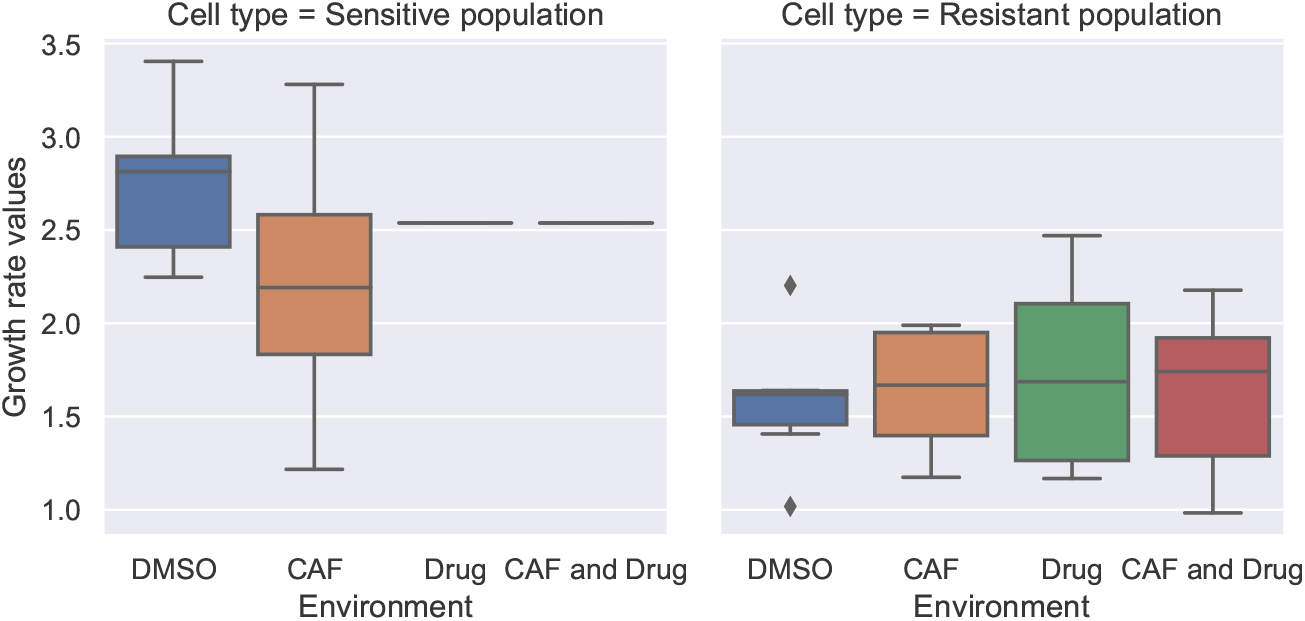
Growth rate parameters (*ρ*_1_ and *ρ*_2_) in all environments. The boxplots of growth rates for sensitive and resistant populations illustrate outcomes in different wells of the monotypic cultures. The growth rate of sensitive cells in the presence of the drug is fixed at the median of those parameters in DMSO and CAF environments. The cell types had significantly different growth rates, while the presence or absence of CAFs and drug resulted in minimal and non-significant variation.

Then, we fixed the sensitive population’s growth rate and carrying capacity parameters at the median of their value in DMSO and CAF environments, to estimate the drug efficacy parameter (*λ*) in environments where Alectinib is present. ANOVA results supported our decision to fix the sensitive population’s growth rate and carrying capacity at the median, showing that population type significantly affected these parameters, while CAFs presence had no significant effect.

We observed no changes in the resistant cells’ population following drug administration in monoculture, observing the time series data. Therefore, we re-estimated the carrying capacity and growth rate parameters for resistant populations in environments containing only the drug and in those with both CAFs and the drug. The boxplots of carrying capacity for sensitive and resistant populations in Fig 3 demonstrate that population type has a more significant influence on the carrying capacity parameter than the presence or absence of CAFs and drug. The lines shown in environments where the drug is present for sensitive cell population correspond to boxplots for fixed values. The variation in carrying capacity estimates of resistant population in the presence of the drug is significantly higher than in other environments, perhaps due to more variations in the initial population in those wells. Although the initial ratio of sensitive and resistant populations is fixed at one and zero, respectively, the initial population size varies in different wells.

Fig 4 illustrates the boxplots of growth rate values. The growth rate estimates for both cancer cell types were not influenced by the environment. We fixed the growth rate of the sensitive population in DMSO and CAF environments and estimated the drug efficacy parameter (*λ*) in environments with Alectinib.

The boxplots for drug efficacy (*λ*), illustrated in Fig 5, show that CAFs significantly decrease the effect of the drug on the sensitive cells’ population. We used the median of (*ρ*_1_, *K*_1_, *ρ*_2_, *K*_2_) parameters in all environments estimated from the monotypic cultures. However, considering that the presence of CAFs significantly influences the drug efficacy term (*λ*), we used the median of this term in the presence and absence of CAFs separately. Using the median of all other parameters, we estimated the competition coefficients (*α*_*SR*_ and *α*_*RS*_) in wells where both populations were present with different initial ratios.

**Fig 5.**
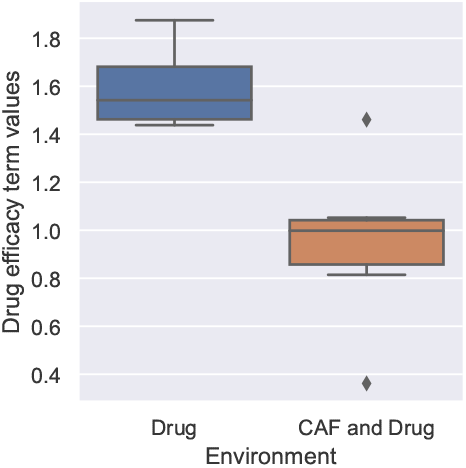
Drug efficacy (*λ*) in environments where drug is present. The boxplots of the drug efficacy parameter for sensitive populations illustrate the fit outcomes in different wells of monotypic culture. The presence of CAFs decreases the effect of the administered drug.

The consideration of carrying capacity, growth rate, and drug effect as varying parameters resulted in overfitting and increased the correlation among the various parameters. Utilizing the estimates derived from the monocultures to construct the two-population model addressed these challenges.

From the boxplots of the competition coefficients derived from six different initial ratios in four environments, illustrated in Fig 6, we see that the presence of the drug increased the effect of sensitive cells on resistant ones through the competition coefficient *α*_*RS*_. However, in the presence of CAFs and drug, the influence of sensitive cells on resistant cells via the competition coefficient *α*_*RS*_ decreases compared to the case where only the drug is administered. Furthermore, the competition coefficients can take values exceeding 1 when the drug is present. In the DMSO environment, the median of *α*_*SR*_ is slightly higher than *α*_*RS*_, which means that the influence of resistant cells on sensitive ones is slightly larger than the other way around. The presence of CAFs slightly decreases the median *α*_*SR*_ and slightly increases the median of *α*_*RS*_. This causes the median of *α*_*RS*_ to become larger than the median of *α*_*SR*_, indicating that CAFs caused the effect of sensitive cells on resistant ones to be higher. However, when the drug is also added to the environment, CAFs have a different effect and decrease the effect of sensitive cells on resistant ones.

**Fig 6.**
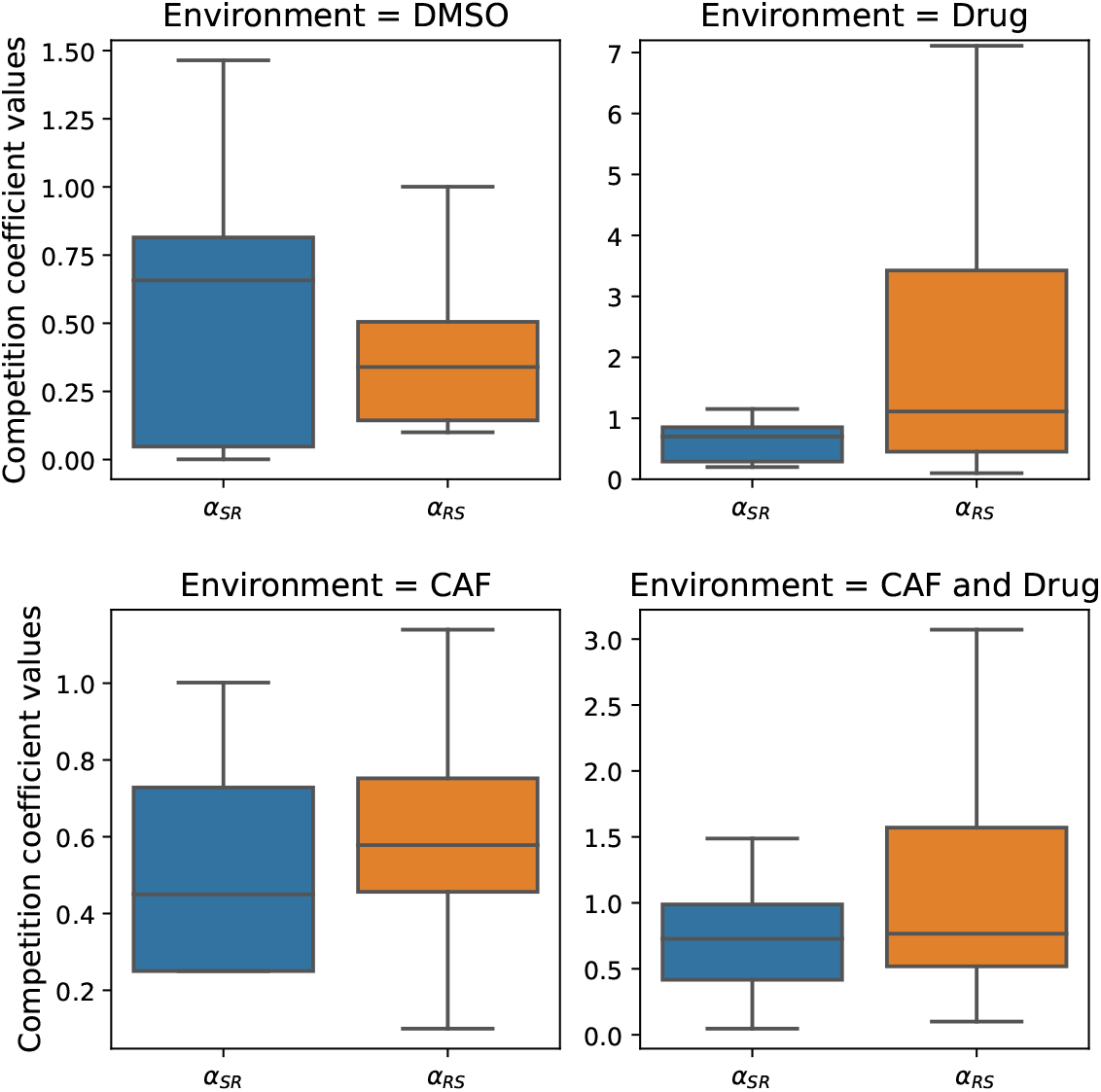
Competition coefficients (*α*_*SR*_ and *α*_*RS*_) derived for ratio-dependent drug effects and a logistic model where all parameters but competition coefficients were fixed. The presence of the drug leads to a large increase in the competition coefficient *α*_*RS*_. Note that the y-axis scale is different in the presence of the drug, and *α*_*RS*_ has values larger than 1.

We used the one-way ANOVA test on *α*_*SR*_ for each pair of environments, which is, in total, six tests. The *α*_*SR*_ values only change significantly when we either add the drug to an environment that has CAFs, or when we add both the CAFs and the drug to the DMSO environment. We repeated the six ANOVA tests for each pair of environments for *α*_*RS*_, and we observed that all environment pairs except the presence of both drug and CAFs, compared to when only the drug is present, led to significant changes. We also performed a one-way ANOVA for *α*_*SR*_ and *α*_*RS*_ in each environment and observed that the effect of sensitive cells on resistant ones, compared to the other way around, is significant in the environment where only the drug is present.

### Environmental influence on competition outcomes between cell types

Considering the median of competition coefficients derived from the co-culture experiment, we examine the outcomes of the competition between sensitive and resistant cell types in different environments by analyzing the Jacobian of the model (Eq 7) and the trajectories. We illustrate the trajectories from three different initial points in Fig 7. In the CAF environment, the model predicts coexistence of the two cell types. In the DMSO environment, the stable equilibrium occurs when the number of sensitive cells is much lower than that of resistant cells. In environments with the drug present, the resistant cells outcompete the sensitive ones.

**Fig 7.**
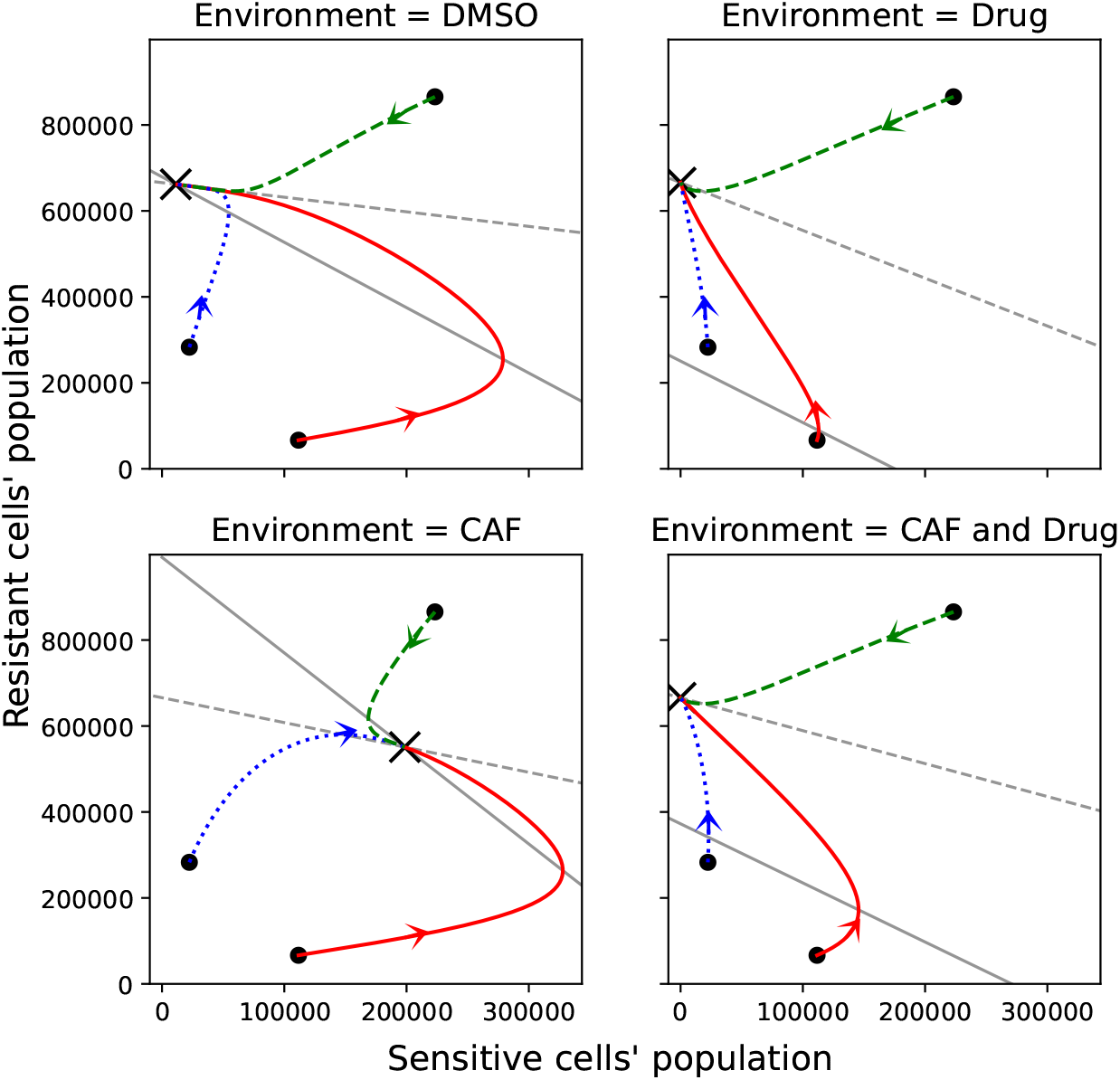
Part of phase plane of the model presented in Eq 7 using the median of the parameters derived from fitting the co-culture data in all environments. The × shows the stable equilibrium in each environment. The equilibrium in the CAF environment shows coexistence of resistant and sensitive cells. The equilibrium in DMSO shows the dominance of resistant cells. The equilibrium in environments where the drug is present shows that sensitive cells become extinct. The trajectories of the model, starting from three randomly chosen initial points, are illustrated in solid red, dashed green, and dotted blue lines. The solid grey line shows 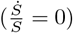 and the dashed grey line shows 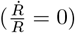, find more detail in

The *ρ*_1_, *ρ*_2_, *K*_1_, and *K*_2_ parameters are considered to be the same in all four environments. Therefore, the different equilibrium properties in CAF compared to those at DMSO is the result of the increase in *α*_*RS*_ and decrease in *α*_*SR*_. The distribution of *α*_*SR*_ is not significantly different in DMSO and CAF (p-value is 0.1). However, the median has a large difference, which strongly influences the equilibrium outcome. The presence of the drug decreases the number of sensitive cells and, through that, changes the equilibrium outcome. For more details on how the change in parameters *λ* and *α*_*SR*_ affects the possibility of mixed equilibria, see Appendix.

## Discussion

Evolutionary game theoretic models can capture cancer eco-evolutionary dynamics and assist in optimizing its treatment [10, 16, 39]. In this paper, we analyzed data from experiments between two cancer cell lines from [25], also analyzed in [40]. We found that the two-population Lotka-Volterra competition model, which incorporates asymmetric competition and ratio-dependent drug efficacy, best captured the observed dynamics and offered novel mechanistic insight into how sensitive and resistant NSCLC cells interact under varying microenvironmental conditions. This ratio-dependent drug effect suggests that treatment effectiveness may depend not only on absolute tumour burden, but also on the relative composition of sensitive and resistant cells — a dynamic feature that can be leveraged in evolutionary therapy design.

Kaznatcheev et al. and Soboleva et al. fitted this data using replicator dynamics and a two-population Gompertz model with Norton-Simon death rate, respectively [25, 40]. They both concluded that these models adequately capture the dynamics of the cancer cells studied. Both studies also reported that Alectinib inhibits the growth of sensitive cells. They also concluded that Alectinib does not affect the growth of resistant cells in monoculture. We confirm these results with our best-fitting model as well.

In the heterotypic culture, Kaznatcheev et al. concluded that cancer-associated fibroblasts (CAFs) enhanced the growth of sensitive cells [25]. We observed that CAFs both decreased resistant cells’ ability to outcompete the sensitive ones and increased the ability of the sensitive cells to outcompete the resistant ones, as seen through median values of the corresponding competition coefficients. While this outcome failed the significance test by a small margin (p-value of 0.1), we believe that if these observations are confirmed in future studies, they can help in designing novel therapies aimed at reducing cancer cells’ competitiveness, for example, by targeting the CAFs concentrations [41–43]. This aligns with growing interest in targeting CAF-mediated signalling to modulate tumour evolution and therapeutic response [41, 42]. Immunotherapy may be the best type of treatment for this purpose [44, 45].

Kaznatcheev et al. observed that resistant cells tend to have higher growth rates than sensitive cells even without the treatment targeting the sensitive cells [25]. Soboleva et al. did not confirm this conclusion [40]. Through our analysis, we found that resistant cells have a larger carrying capacity compared to sensitive ones, rather than having larger intrinsic growth rates. This distinction is critical for designing therapies that constrain tumour expansion without necessarily suppressing proliferation — an approach already shown to be effective in prostate cancer [16, 46] and potentially translatable to NSCLC. This nuance could not be observed with the simpler models of Kaznatcheev et al. and Soboleva et al. As the first evolutionary therapies in prostate cancer targeted the carrying capacity of cancer cells rather than their growth rate [46], this suggests opportunities for optimizing treatment non-small cell lung cancer as well.

Kaznatcheev et al. concluded that in the environment where only CAFs are present, the sensitive and resistant cells can coexist while in all other environments, the resistant cancer cell population will outcompete the sensitive population [25]. We also conclude that coexistence of the twopopulations is possible when only CAFs are present in the environment. In contrast, in environments where the drug is present, it leads to a fully resistant, stable equilibrium point. However, our model also predicts the coexistence of sensitive and resistant cells in DMSO, albeit with low abundances of sensitive cells.

Kaznatcheev et al. concluded that resistant cells will outcompete sensitive cells under DMSO. Furthermore, the drug efficacy parameter *λ* is smaller in the presence of both CAFs and the drug compared to the environment where only the drug is present. We also confirm the positive effect of cancer-associated fibroblasts on the sensitive cells relative to the resistant cells.

Our model considers two types of cancer cells, one resistant to the Alectinib drug and one sensitive to it, that are well-mixed in the environment. While our model assumes a well-mixed population, this may not fully reflect spatial dynamics present in the experimental setup. If the two populations are not well-mixed, spatially explicit models may be more suitable to model their dynamics [47]. We could use agent-based, partial-differential equation, or other types of spatial models to address this issue [19, 48, 49]. Furthermore, we do not consider the resistance level of resistant cells as an evolving trait [14, 50, 51]. This aspect could be an interesting aspect for future research, as the resistant cells may evolve in the course of the experiments if they last sufficiently long.

We estimated how the presence of CAFs and the Alectinib drug affect the model parameters and, through that, the outcome of competition between sensitive and resistant cells. We presented how the presence of CAFs results in the coexistence of sensitive and resistant cells. Reaching the studied equilibria may be possible through dose adjustment in response to the anticipated tumor burden in [14, 51–53].

Experiments are usually not performed to analyze both steady state behavior of the studied system and the transient dynamics leading towards this behavior. Even if that is the case, the number of replicates is often limited due to practical constraints. Here, we utilized our prior knowledge of the cells’ behavior and estimated model parameters for analyzing the steady-state behavior of cancer cells under different conditions.

The model we have found to be the most appropriate for capturing the dynamics of NSCLC within these experiments could form the basis for model-informed treatment. Of interest is treatment optimization within this model, to explore and compare treatment strategies to improve patients’ quality of life and survival. This aligns with the systems perspective emphasized by Soboleva et al. [54], where interdisciplinary models inform adaptive treatment strategies aimed at improving patient outcomes. As cancer therapy moves toward dynamic, model-informed approaches, frameworks like ours will be essential for translating eco-evolutionary insights into clinically actionable strategies.

## Acknowledgments

We thank Dr. Artem Kaznatcheev for making the in-vitro data from [25] available, which enabled this research. We are also grateful for his insightful comments and clarifications, which deepened our understanding of the dataset and contributed to the improvement of the manuscript.

## S1 Appendix

**General form of two-population models with Gompertz and Von Bertalanffy growth**The general form of two-population models with Gompertz and Von Bertalanffy growth and Norton-Simon drug effect is presented in Tables 2, 3.

## S2 Appendix

**AIC goodness of fit measure** Fig 8 illustrates the AIC value of the analyzed 15 models.

**Fig 8.**
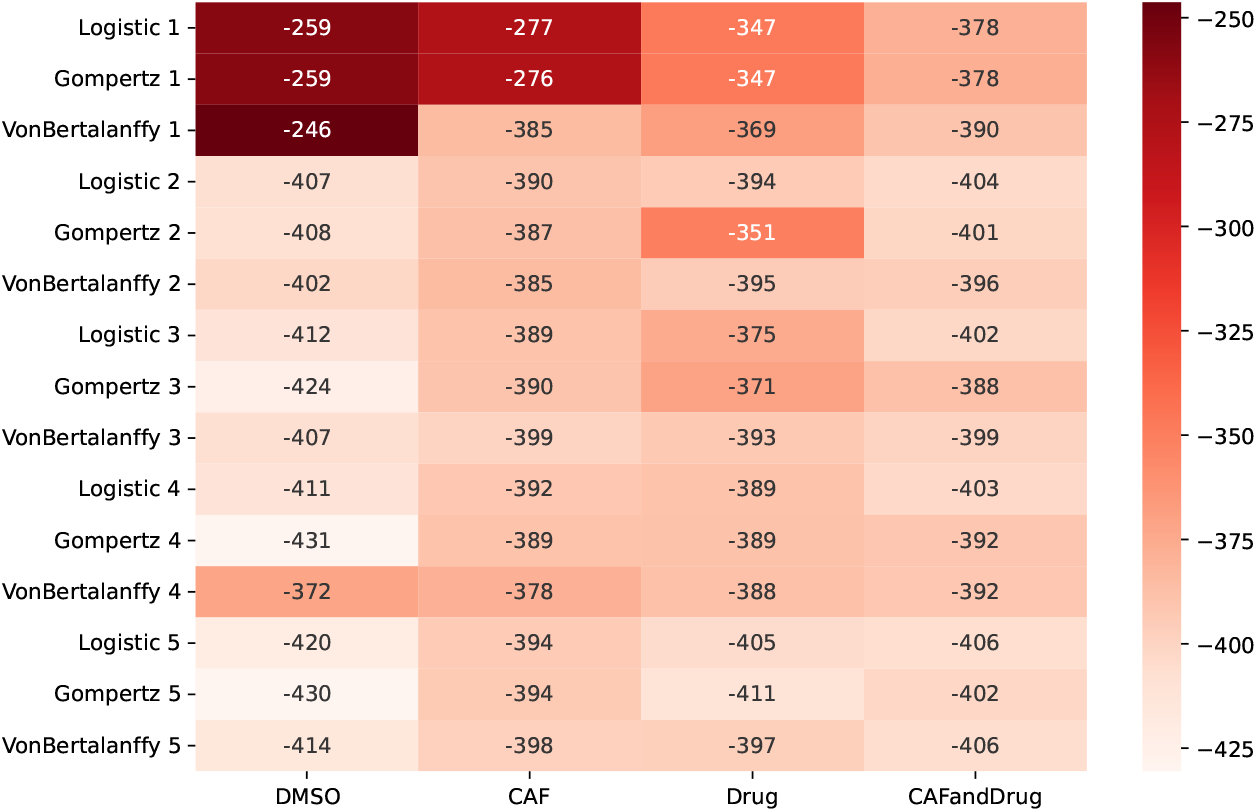
AIC result of the two-population model fits. Heatmap of the fit results for the proposed fifteen models in Tables 1, 2, 3. Models with more than two unknown parameters in drug-free environments and more than three parameters when the drug is present fit the data.

## S3 Appendix

**ANCOVA test on relative growth** To determine the growth dynamics, we analyze the relation between the population’s relative growth 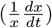 with respect to the population (*x*). By determining the nonlinearity order of the relation, we can differentiate between logistic, Gompertz, and Von Bertalanffy growth dynamics.

The analysis is done for sensitive and resistant cells within monotypic cultures in DMSO and CAF environments. We employ the ANCOVA test to identify the factors that significantly influence the relative growth of the cell population. The specific variables examined include the number of the well, the cell population, their cross-correlation, the intercept, and the second-order term of the population. The selected factors hold significant biological and experimental relevance to growth dynamics: the number of wells and cell population serve as primary experimental variables, cross-correlation addresses potential interactions, the intercept reflects baseline effects, and the second-order term allows the examination of nonlinear relationships in growth behavior. First, we analyze the relative growth explainability with cell population as the continuous variable, well number as the categorical variable, and their correlation 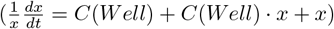. We observe that the population’s and intercept’s effects are statistically significant in DMSO and CAF environments since the p-values of the population variable and intercept for sensitive and resistant cells are less than 0.001. In the next step, we analyzed the significance of the second order of the population variable. For the model with the first order of population, the second order of population, and intercept on the right-hand side 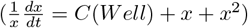, the second-order term was not statistically significant for sensitive and resistant populations. We conclude that the Logistic model is more suitable here than Gompertz and Von Bertalanffy since the relative growth is linearly related to the population.

## S4 Appendix

**Nullcline of the population model**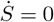 result in two lines *S* = 0 and 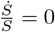 and 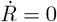 result in two lines *R* = 0 and 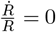. The intersection of *S* = 0 and 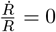 results in equilibrium point *P*_1_ at (*S, R*) = (0, *K*_2_). The intersection of *R* = 0 and 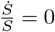 results in equilibrium point *P* at 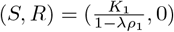. The intersection of 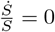 and 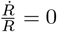 results in a mixed equilibrium point, *P*_3_, which might not exist for some parameter values. The value of 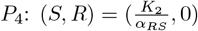 and 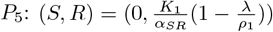 could be higher and lower than *P*_2_ and *P*_1_, respectively, depending on parameter values. This determines whether the mixed equilibrium point exists or not. As illustrated in Fig 9, depending on *α*_*SR*_, *α*_*RS*_, and *λ* values, there can be cases where the mixed equilibrium located at *P*_3_ does not exist. For our parameter values, *P*_4_ is always larger than *P*_2_. However, in the presence of the drug, *λ* and *α*_*SR*_ become larger. Due to this change, *P*_5_ becomes smaller than *P*_1_, which leads to disappearance of the mixed equilibrium points and extinction of sensitive cells. Furthermore, adding only CAFs to the DMSO environment leads to a decrease in *α*_*SR*_, which causes *P*_5_ to be larger than *P*_1_, leading to a mixed equilibrium point with a large population of sensitive cells.

**Fig 9.**
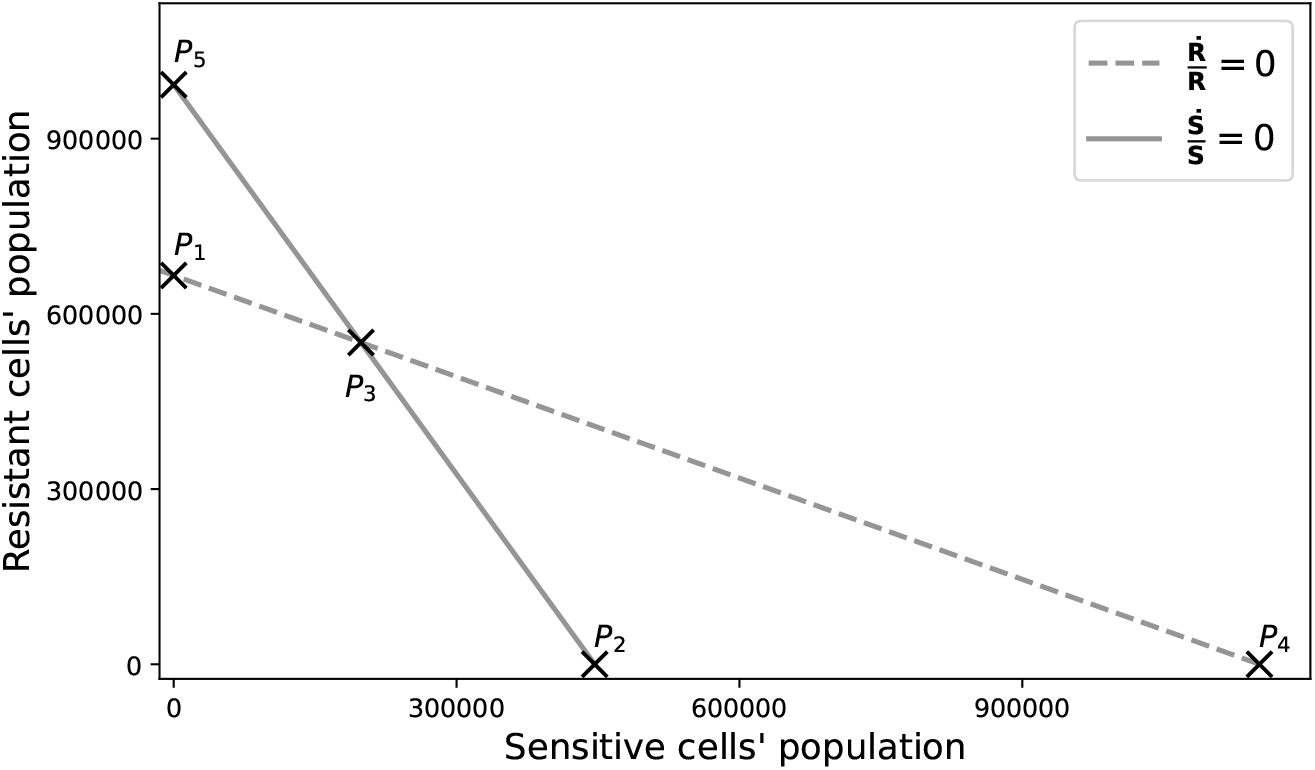
Intersection of nullclines of Eq 7. Setting the derivative of sensitive population to zero, 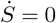, results in two lines *S* = 0 and 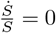. Setting the derivative of resistant population to zero, 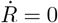, results in two lines *R* = 0 and 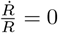 point shows (*S, R*) = (0, *K*2); *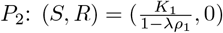*; *P*3 shows the mixed equilibrium which might not exist for some parameter values; 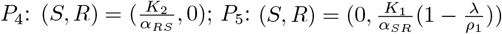.

